# Cellular advective-diffusion drives the emergence of bacterial surface colonization patterns and heterogeneity

**DOI:** 10.1101/434167

**Authors:** Tamara Rossy, Carey D. Nadell, Alexandre Persat

**Affiliations:** Institute of Bioengineering and Global Health Institute, School of Life Sciences, École Polytechnique Fédérale de Lausanne, Lausanne, Switzerland; Department of Biological Sciences, Dartmouth, Hanover, NH 03755, USA

## Abstract

Microorganisms navigate and divide on surfaces to form multicellular structures called biofilms, the most widespread survival strategy found in the bacterial world^1–4^. One common assumption is that cellular components guide the spatial architecture and arrangement of multiple species in a biofilm. However, bacteria must contend with mechanical forces generated through contact with surfaces under fluid flow, whose contributions to colonization patterns are poorly understood. Here, we show how the balance between motility and flow promotes the emergence of morphological patterns in *Caulobacter crescentus* biofilms^5^. By modeling transport of single cells by flow and Brownian-like swimming, we show that the emergence of these patterns is guided by an effective Péclet number. By analogy with transport phenomena we show that, counter-intuitively, fluid flow represses mixing of distinct clonal lineages, thereby affecting the interaction landscapes between biofilm-dwelling bacteria. This demonstrates that hydrodynamics influence species interaction and evolution within surface-associated communities.

---

Bacterial motility plays a key role during early biofilm growth^5–8^, but the joint roles of fluid flow and bacterial surface interactions have only just begun to receive attention^9^. As one might suspect, hydrodynamic forces can disrupt biofilms^10^, but less intuitively, they can also promote unusual biofilm structures^11^ or alter the evolutionary dynamics of matrix secretion^12^. Given the importance of fluid flow in remodeling biofilms and in transporting planktonic cells or aggregates, we anticipate that such forces also modulate spatial organization of surface associated bacterial collectives on many scales.

Most microbes have evolved cellular components optimizing their interactions with surfaces^2,13,14^. *C. crescentus* is particularly well-adapted to life on surfaces under flow: polar stalk and holdfast confer strong attachment and its curved morphology promotes biofilm formation in flow^15^. During the process of growth on surface, *C. crescentus* mother cells asymmetrically divide into a non-motile stalked cell and a new motile daughter swarmer cell^16^. The characteristic curved shape of *C. crescentus* promotes local surface colonization of daughter cells by reorienting the body of sessile mother cells in the direction of the flow, accelerating accumulation of biomass near the founder cell^17^. Thus, during sessile division in flow, daughter swarmer cells may either attach immediately downstream their mother or explore the surrounding fluid to later reattach. The former depends on cell shape while the latter must depend on fluid transport mechanisms. The relative importance of these two surface colonization modes will, we predict, dramatically influence the basal architecture and cell lineage structure of nascent biofilm populations.

To investigate the contributions of fluid flow to surface colonization patterns, we exposed surface-associated cells to controlled flow in microchannels. In relatively weak flow (2 mm.s^−1^), *C. crescentus* rapidly and uniformly colonizes the surface (Fig. 1a, top). In contrast, spatial patterns of colonization emerged in strong flow (27 mm.s^−1^), where biofilms grew into sparse, dense microcolonies (Fig. 1a, bottom). Surface occupation dramatically drops for growth at mean fluid velocity higher than 4 mm.s^−1^ (Fig. 1c, left). Visualization at higher spatial resolution highlights the presence of many isolated single cells in weaker flow, which are absent in stronger flows. As a result, clusters are generally small in weak flow (< 40 μm^2^), in comparison to strong flows (>100 μm^2^) (Fig. 1c, right). Thus, flow promotes the emergence of multicellular patch-like patterns at the channel surface but slows down surface occupation.

**Figure 1:**
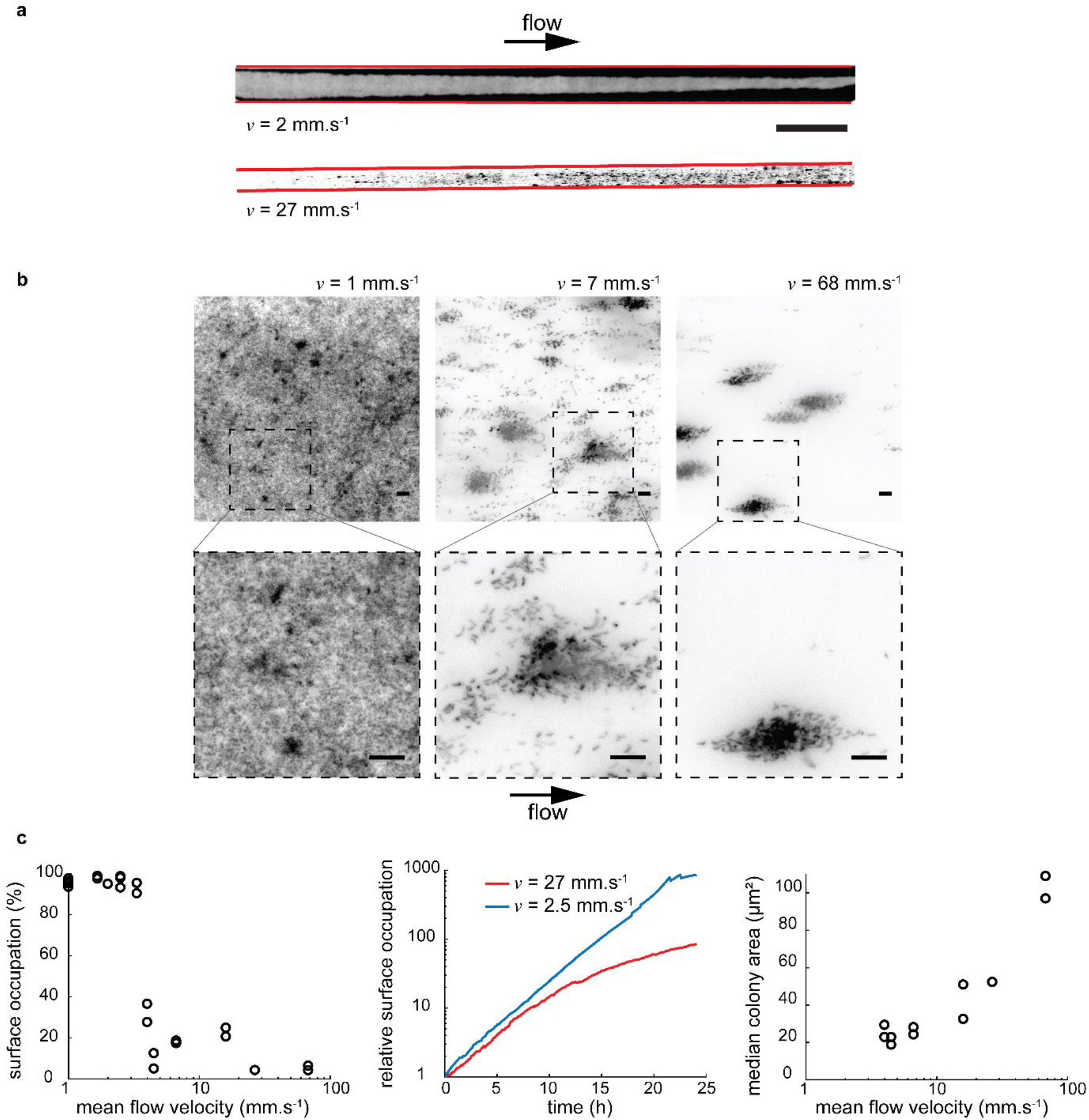
Flow modulates *C. crescentus* colonization patterns. **(a)** Grayscale display of fluorescence microscopy images of *C. crescentus* after 48h exposure to fluid flow in microchannels. In weak flow (top), surface colonization is uniform. In strong flow (bottom), biofilms grow into patterns of discrete cell clusters. The edges of the microchannel are highlighted in red. Scale bar: 1 mm. **(b)** Colonization patterns at the channel centerline at three representative flow velocities, after 24h of colonization under flow. The bottom images show close-up views to distinguish single cells. In weaker flow (left), the channel surface is nearly saturated. At intermediate flow (middle), multicellular clusters are surrounded by smaller groups or single isolated cells. In strong flow (right), biofilms grow mainly as multicellular clusters. Scale bars: 10 μm. **(c)** Fluid flow modulates kinetics and pattern geometry during surface colonization. Surface occupation after 24h of growth as a functionof mean flow velocity (left). Each data point corresponds to an individual experiment. Surface occupation over time for two representative flow velocities (center). Median microcolony area after 24h of growth as a function of mean flow velocity (right).

During *C. crescentus* asymmetric division, unattached progeny is released into the fluid bulk (Fig. 2a). Dynamic visualizations of surface colonization highlight that, in strong flow, biofilms develop from single founder cells (Supplementary Movie 1), whereas in weak flow, new founder cells frequently attach to the surface, speeding up the overall rate of colonization and homogenizing surface coverage (Supplementary Movie 2 and Fig. 1c, center). We thus suspected that the relative contribution of random spatial exploration by swimming motility, and flow transport may enable reattachment. To demonstrate this, we abolished swimming motility by deleting the flagellar gene *flgE.* For this strain, flow is the dominant transport mechanism of swarmer cells. Fig. 2b shows a comparison between biofilms formed by wild-type (WT) and *flgE*^−^ in weak flow. In contrast to WT, flagellum-less cells colonize the surface into patch-like patterns and with an apparent decrease in single isolated cells, similarly to the WT in intermediate flow (Fig. 1b, center). This demonstrates that motility plays a critical role in controlling surface occupation density and distribution.

**Figure 2:**
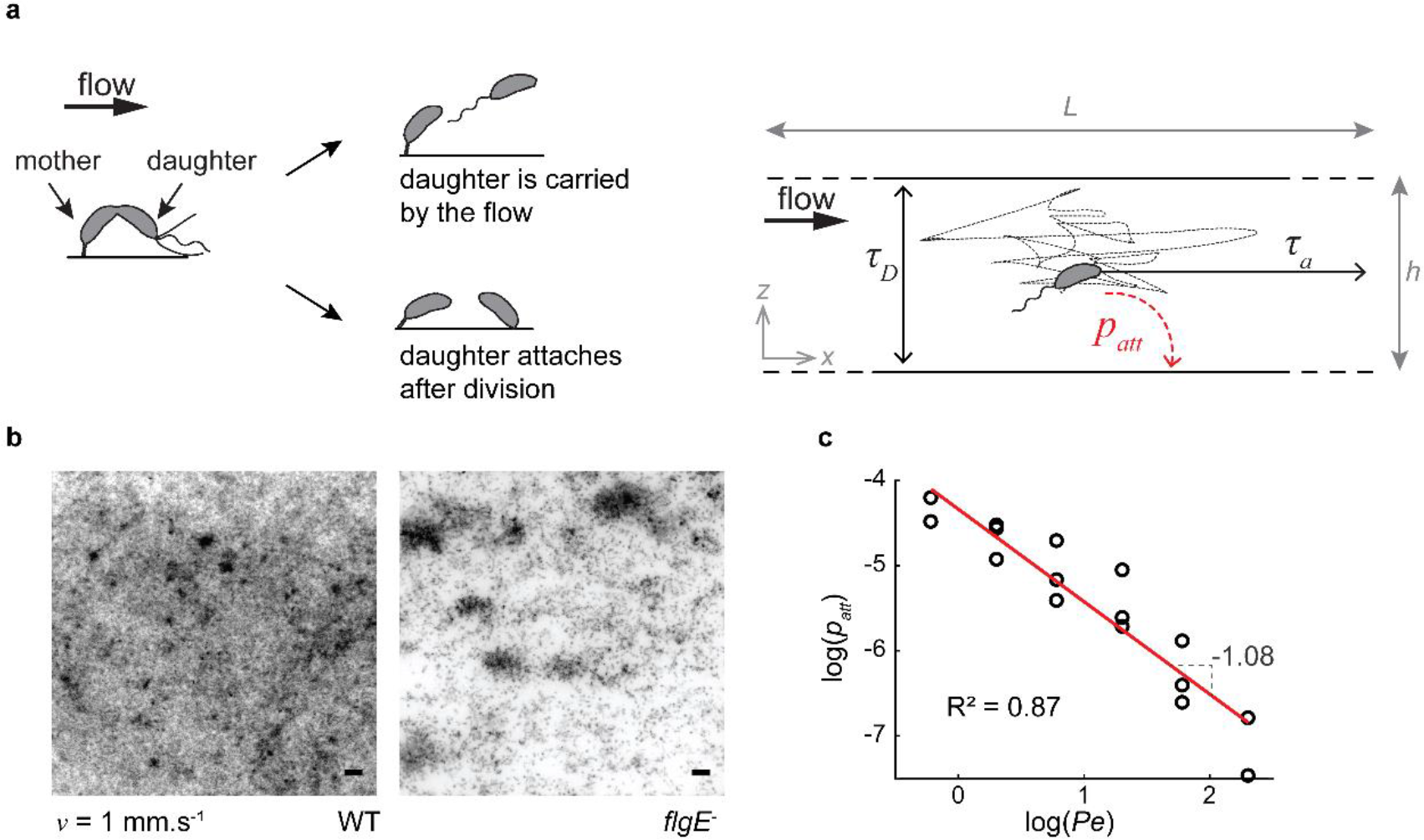
Physical mechanism for modulation of *C. crescentus* biofilm architecture. **(a)** *C. crescentus* divides asymmetrically: the mother cell is anchored to the surface and undergoing division. At the time of division, a daughter cell can either attach to the surface or be carried by the flow. If attachment occurs, the daughter immediately synthesizes a holdfast contributing to clonal expansion on the surface. If the daughter cell does not attach to the surface, it is subject to: (i) advective transport by fluid flow and (ii) diffusion-like transport generated by unbiased swimming. **(b)** Contribution of bacterial motility to surface colonization patterns. Fluorescence microscopy images of wild-type (WT) or flagellum-less (*flgE*^−^) *C. crescentus* after 24h exposure to fluid flow (1 mm.s^−1^). *flgE^−^* colonizes the surface less densely and less uniformly than WT, qualitatively recapitulating the results observed in stronger flow for WT. Scale bars: 10 μm. **(c)** Attachment probability (attachment rate normalized by total bacterial flux) as a function of the Peclet number (*Pe*) on a logarithmic scale. A linear fit of the data indicates swarmer adhesion probability scales with Pe-^1^, as suggested by our advective-diffusion model.

Flow transports bacteria directionally along streamlines, whereas cells swim in diffusive, Brownian-like trajectories in the absence of chemical gradients^18^. We therefore drew an analogy with advective-diffusion transport problems: the balance between flow-driven advective transport of single cells and their diffusive flagellar motility must contribute to the distinct colonization patterns observed in our experiments. We thus developed a scaling for the probability of attachment of a free-swimming bacterium as a function of fluid velocity by reasoning in terms of timescales. Fluid flow transports swarmer bacteria from their division site towards the channel outlet in a characteristic time *τ*_*a*_ = *L*/*ν*, where *L* is the microchannel length and *v* the mean flow velocity. During this time, a cell explores the depth of the channel by swimming, effectively diffusing in the direction perpendicular to the surface with characteristic timescale *τ*_D_ = *h*^2^/*D*, where *h* is the channel height and *D* the effective diffusion coefficient of a bacterium attributed to unbiased swimming^18^. The probability that a free-swimming bacterium reattaches to the surface depends on the ratio of these two timescales: *τ*_D_/*τ*_a_ = (*h*^2^*ν*)/(*DL*), a non-dimensional quantity resembling a Péclet number (*Pe*), which measures the relative contributions of advective to diffusive transport^19^. At large *Pe* (*τ*_*a*_ ≪ *τ*_*D*_) cells are rapidly washed out of the channel before encountering the surface so that the probability of attachment is low (*p*_*att*_~0). In contrast, at very low *Pe* diffusion dominates over flow; a planktonic cell has sufficient time to reach the surface before being flushed out of the channel, and may eventually reattach to the surface away from its stalked parent (*p*_*att*_~1). To validate this scaling, we measured the attachment rate of planktonic cells as a function of applied flow velocity. We counted the number of cells attaching onto the surface per unit time, and estimated the corresponding *p*_*att*_. We found that attachment probability scales with Pe^−1^ (Fig. 2c), which is consistent with the advection-diffusion model, validating our physical explanation of surface colonization rates.

While patterns of surface colonization are crucial for initiating biofilm growth, they can also set the foundation for clonal lineage structure, a key factor influencing the evolution of microbial interaction traits^20^. The spatial structure of biofilms is heterogeneous and dynamic^20^. Natural biofilms are thought to commonly include multiple strains and species that can be organized in a variety of threedimensional patterns^21^. The spatial arrangements of such multi-species consortia can dramatically impact evolution of cell-cell interactions, and vice versa^20^. The mechanisms by which environmental conditions, such as fluid flow, and microbial response to these factors influence the spatial architecture of polymicrobial communities, however, are still unclear.

In mass transport phenomena, the balance between advective and diffusive transport strongly influences mixing of fluids and solutes^19^. By analogy, we reasoned that since surface occupation by *C. crescentus* is governed by advective diffusion, flow may also impact the mixing of distinct cell lineages and their social interactions. We grew *C. crescentus* biofilms in various flow conditions, starting from a one-to-one mixture of strains constitutively expressing mKate or Venus fluorescent proteins whose doubling times are identical^17^ (Fig. 3a). Consistent with advective-diffusion transport, surface populations of mKate- and Venus-expressing cells were well mixed in weak flow. There was no clear region where clonal lineages were segregated at scales larger than 10 μm. Within seemingly homogeneous clonal groups of cells, we could generally find invaders expressing the other fluorescence protein. At higher flow velocity, clonal groups were larger and segregated from each other, suggesting that they originated from a single parent cell.

**Figure 3:**
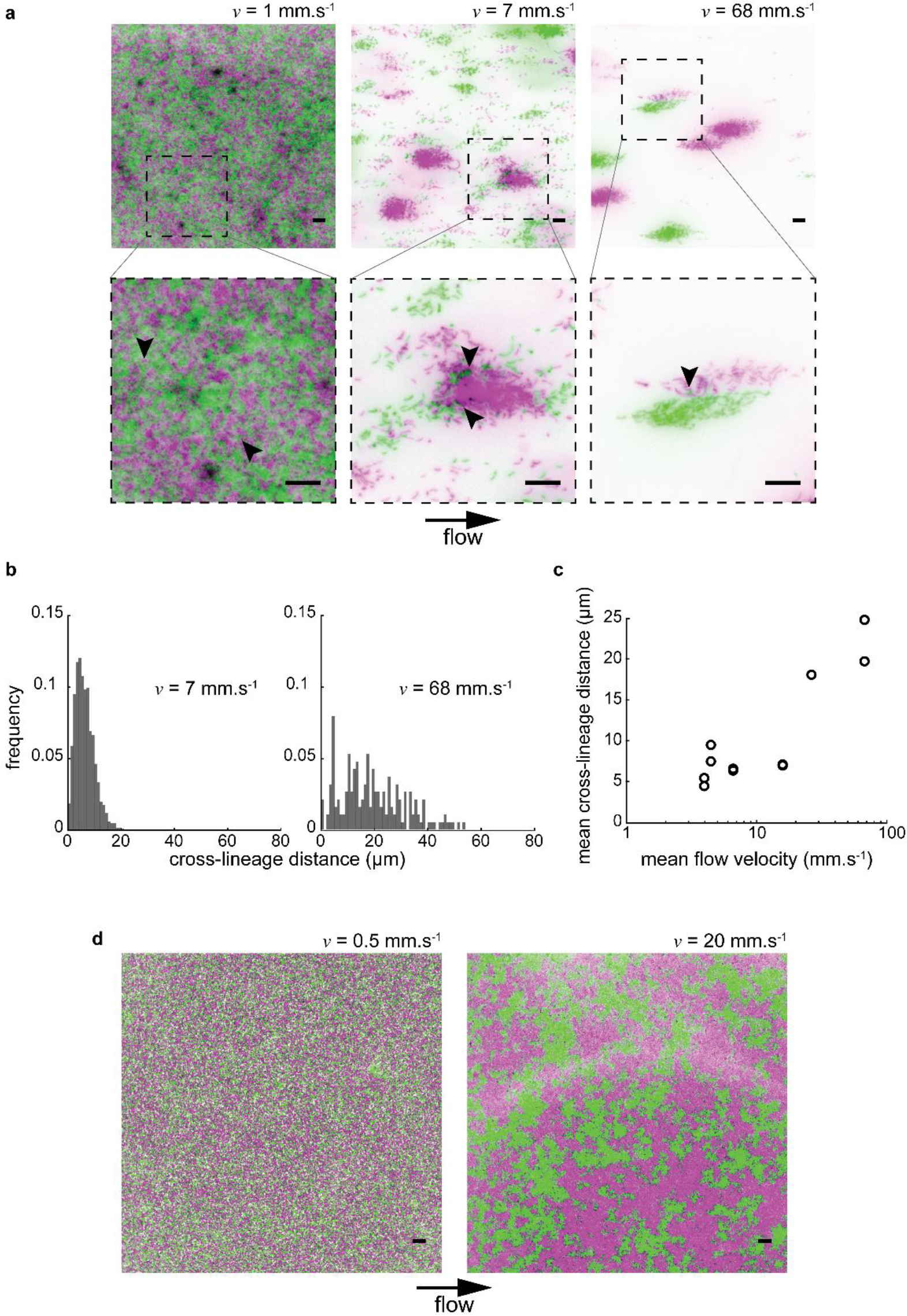
Flow modulates clonal structuring of *C. crescentus* biofilms. **(a)** Fluorescence microscopy images of *C. crescentus* biofilms (24h). Two populations at equal density, expressing either mKate or Venus fluorescent proteins, were initially loaded in microchannels. The bottom row of images highlights the presence of invading cells (indicated by black arrowheads) within otherwise clonal clusters. Green: *C. crescentus* mKate. Magenta: *C. crescentus* Venus. Scale bars: 10 μm. **(b)** Distribution of cross-lineage colony distances (i.e. distance between green colonies and their nearest magenta neighbor, and vice-versa) for two representative mean flow velocities (7 mm.s^−1^ and 68 mm.s^−1^). The distribution broadens as flow velocity increases. **(c)** Crosslineage colony distance, which can be used as a measure of clonal segregation, as a function of mean flow velocity. As flow velocity increases, the mean cross-lineage distance increases, indicating that biofilm mixing decreases. **(d)** *C. crescentus* biofilms on the surface of a microchannel after 6 days of growth under flow. They recapitulate the initial patterns of colonization shown above. Scale bars: 10 μm.

The distribution of cross-lineage colony distances effectively measures segregation and thus strongly depends on flow intensity: in intermediate flow, all colonies expressing a given fluorescent protein are at most ~20 μm away from their nearest counterpart (Fig. 3b). The distribution is heavily weighted at low values of nearest neighbor distance (standard deviation = 3.5 μm). In contrast, at high flow intensity, the distribution of cross-lineage colony distances broadens dramatically (standard deviation = 12.8 μm). Colonies from each color variant can be separated by as much as 50 μm, and there is a substantial decrease in the frequency of small intercolony distances. This shift in distribution occurs progressively as flow intensity increases: the mean cross-lineage distance indeed increases as a function of mean flow velocity, demonstrating that segregation strengthens with flow (Fig. 3c). These observations are consistent with a model where the balance between advection and diffusion of planktonic cells and deposition of daughter cells adjacent to their points of origin dictates the level of clonal structure within nascent *C. crescentus* biofilms. At high *Pe*, flow represses mixing of clones by carrying planktonic cells far from their parent, while at low *Pe* motility drives diffusive swimming trajectories to promote clonal mixing (Supplementary Fig. 1). We confirmed this by observing a reduction of clonal mixing of flagellum-less mutant at low flow intensity (Supplementary Fig. 2). Finally, we noted that clonal patterns are conserved later in the colonization process. After 6 days of growth, biofilms in both weak and strong flow regimes covered the surface entirely and extended into the channel depth, but, importantly, retained the clonal structure set by the initial patterns of surface occupation (Fig. 3d). The three-dimensional spatial distribution of clones remained highly mixed at low flow and relatively segregated at high flow (Supplementary Fig. 3).

We demonstrated that the multi-scale feedbacks between surface attachment, daughter cell deposition, fluid transport, and dispersion by diffusion exert a strong influence on the morphological, spatial and genetic structure of biofilm populations. The early stages of surface colonization can set the foundations of subsequent biofilm architecture, influencing the spatial distributions of different strains and species, and the community’s interaction networks. One critical ingredient to this process is probabilistic local attachment versus planktonic release of daughter cells. Such asymmetries in adhesive properties may very likely appear between two daughter cells that are dividing symmetrically^22^. For example, a memory effect in *Pseudomonas aeruginosa* yields strong differences in the adhesive behavior of two sessile daughter cells, nearly recapitulating the pattern of *C. crescentus*, despite the absence of obvious cellular asymmetry^6,23,24^. The balance between directional advective and random diffusive trajectories of planktonic cells constitutes a second ingredient setting the spatial structure of biofilm communities. A Péclet number can be used to predict the emergence of motility- and flow-induced morphological transitions. In the same manner, a Péclet number quantifying the relative contribution of directional cell displacement to rotational diffusion describes phase-like transitions between multicellular phases in *Myxococcus xanthus*^25^.

## Author Contributions

T.R., C.N. and A.P conceptualized the study, T.R. performed experiments and data analysis. T.R., C.N. and A.P. wrote the manuscript.

## Acknowledgements

TR and AP are supported by the Swiss National Science Foundation, Projects grant 31003A_169377 and the Giorgio Cavaglieri Foundation. CDN is supported by the National Science Foundation (MCB 1817342), a Burke Award from Dartmouth College, a pilot award from the Cystic Fibrosis Foundation (STANTO15RO), and NIH grant P20-GM113132 to the Dartmouth BioMT COBRE.

## Data availability

All data are available from the corresponding author upon reasonable request.

## Code availability

All codes are available from the corresponding author upon reasonable request.

## Methods

### Design and fabrication of the microfluidic chips

We fabricated the microfluidic chips following standard soft lithography techniques. More specifically, for the 24h- and 48h-long biofilm experiments, we designed 1 cm-long, 500 or 250 μm-wide channels in Autodesk AutoCAD 2018 and printed them on a soft plastic photomask. We then coated silicon wafers with photoresist (SU8 2025, Microchem), with varying thicknesses (25 μm, 50 μm and 90 μm) to allow a wider range of mean flow velocities for identical flow rate settings. The wafer was exposed to UV light through the mask and developed in PGMEA (Sigma-Aldrich) in order to produce a mold. PDMS (Sylgard 184, Dow Corning) was subsequently casted on the mold and cured at 80°C for about 1h30. After cutting out the chips, we punched 1 mm inlet and outlet ports. We finally bonded the PDMS chips to glass coverslips (Marienfeld 1.5) in a ZEPTO plasma cleaner (Diener electronic). To fabricate channels for the 6 day-long biofilm experiments, we followed a similar procedure, but adjusted the dimensions of the channel to leave more space for large 3D structures to form. More precisely, the channel was 2 mm wide, 110 μm high.

### Bacterial strains

We used strain CB15 constitutively expressing a fluorescent protein (Venus or mKate)^17^. These bacteria grew in peptone yeast extract medium supplemented with 5 μg/ml of kanamycin (PYE-Kan), in a shaking incubator set to 30°C. For the experiments involving non-motile CB15, we inserted either mKate or Venus in a flagellum-less mutant, CB15 *flgE-* ^17^. We prepared electrocompetent CB15 *flgE-* by centrifuging 3 ml of stationary phase culture and rinsing it two times with cold Milli-Q water (Merck Millipore). About 600 ng of plasmid (either Venus or mKate) were added for transformation and the bacteria were then plated on PYE-Kan plates.

### Biofilm growth in microfluidic chambers

At the start of every experiment, the bacterial cultures had an optical density of approximately 0.15 (~ 4.5·10^8^ CFU.ml^−1^). Equal volumes of CB15 mKate and CB15 Venus were diluted in PYE-Kan to a final 1:10 concentration. We then loaded the bacterial mixture in a microchannel using a micropipette, and let them adhere for 3 minutes before washing the channel with PYE-Kan. For all conditions but the highest flow velocity (*v* = 68 mm.s^−1^ for 24h biofilms, and *v* = 20 mm.s^−1^ for 6-day biofilms), we connected the inlet port to a disposable PYE-Kan-filled syringe (BD Plastipak) using a 1.09 mm outer diameter polyethylene tube (Instech) and a 27G needle (Instech). The syringe was then mounted onto a syringe pump (ZS100, ChuangRui Pump). For the highest flow conditions (*v* = 68 mm.s^−1^ for 24h biofilms, and *v* = 20 mm.s^−1^ for 6-day biofilms), we connected the inlet port to a PYE-Kan-filled beaker via two imbricated tubes (polyethylene tubing as described above, and Tygon-LFL tubes with an inner diameter of 0.76 mm (Ismatec)). We mounted the setup onto a peristaltic pump (MCP, Ismatec) allowing us to work with larger volumes than the syringe pump. For every experiment, we connected the outlet port to a waste container using polyethylene tubing. We finally placed the chip in a 30°C incubator and applied a controlled flow of PYE-Kan to the microchannels for 24h, 48h or 6 days depending on the experiment. The mean flow velocity (v) was calculated from the selected flow rate (Q) and channel cross-sectional area (A) as such: *ν* = *Q/A*.

### Visualization

For all visualizations of biofilms grown up to 48h, we used a Nikon TiE epifluorescence microscope equipped with a Hamamatsu ORCA Flash 4 camera and a 40X Plan APO NA 0.9 objective. The full-channel images were stitched using the NIS-Elements software. All single cell level pictures presented in this work were taken 9 mm away downstream of the inlet. For the timelapse experiments (Supplementary Movies 1 and 2), we acquired images every 5 minutes for 24 hours. To visualize 6 day old biofilms, we used a Leica SP8 confocal microscope equipped with a white laser, a 25X HC FLUOTAR NA 0.95, water-immersion objective, as well as a 63X HC PL APO NA 1.40 oil-immersion objective for high magnification z-stack acquisitions. We used Imaris (Bitplane) for three-dimensional rendering of z-stack pictures (Supplementary Fig. 3).

### Data analysis

Data analysis was conducted using Matlab (Mathworks). The images were segmented using an adaptive threshold, the sensitivity of which varied depending on the median intensity of the picture. Similarly, the percentage of background removed was also determined by the median intensity. Finally, we filtered out objects smaller than 15 pixels, since this value was observed to be the minimal area of a single cell standing vertically. After segmentation, pictures were visually assessed to ensure good quality of segmentation. In rare cases (4 pictures out of 50), segmentation was unsatisfying and the images had to be excluded from the analysis.

To calculate the surface coverage and microcolony area, we merged the segmented pictures originating from mKate and Venus using the logical *or* function. To quantify surface coverage, we divided the area of white pixels (i.e. pixels containing a part of cell) by the total area of an image.

We observed that single cell clusters were difficult or even impossible to discriminate by eye when surface coverage was larger than 80%. Therefore, we only included segmented pictures with a surface coverage ≤ 80% for the measurement of microcolony area. We also filtered out any object smaller than 200 pixels, which approximately corresponds to a group of five cells (average cell size: 1.29 μm^2^ ≈ 40.2 pixels, N = 80 cells). We then closed the pictures using a built-in Matlab function (with a disk structuring element having a radius of five pixels) and calculated the area of every colony. The median colony area was finally calculated for each image.

To quantify the degree of mixing of the biofilms, we again only studied segmented pictures with a surface coverage ≤ 80%. Additionally, unlike for surface coverage and colony area quantification, we analyzed mKate and Venus pictures separately. We closed all the pictures as mentioned above. We then calculated the distance between the centroid of an object and its nearest neighbor expressing the other fluorescent protein, using the built-in Matlab function *knnsearch.* This operation was repeated for every object in every picture. Finally, the mean cross-lineage distance was calculated for each experimental condition, taking into account distances from both fluorescently-labeled populations.

### Estimation of the probability of attachment as a function of Pe

To estimate the attachment probability of swarmer *C. crescentus* in different flow conditions, we flowed CB15 Venus cells in a 500 μm-wide, 90 μm-high microchannel using a syringe pump. The flow rates varied between 0.81 and 270 μl.min^−1^ (mean flow velocities from 0.3 to 100 mm.s^−1^ respectively). Each condition was repeated two to three times. Bacteria were visualized by fluorescence microscopy (one frame recorded every second during one minute) and single attachment events were counted. Bacteria had to remain on the surface for at least 3 consecutive frames at the same location to be counted as attached. The number of bacteria attached over time was plotted for each flow condition and, using a linear fit, we extracted the attachment rate from the slope of these curves. The attachment probability was then computed as follows:

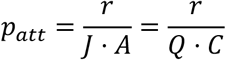

Where *r* is the attachment rate, *C* is the bacterial concentration and *J* is the bacterial flux, defined as *J* = (*Q* · *C*)/*A*. Note that the bacteria loaded in the channel contained a mixture of swarmer and stalked cells, thus our measurement of *p*_*att*_ is underestimated compared to biofilm growth conditions where all planktonic cells are swarmers.

To determine the dependence of attachment on the flow regime, we plotted the logarithm of *p*_*att*_ as a function of the logarithm of *Pe*. As defined above, *Pe* = (*h*^2^*ν*)/(*DL*); we assumed the diffusion coefficient of *C. crescentus* to be equal to that of *E. coli* reported before^18^, namely *D* = 4·10^−6^ cm^2^.s^−1^. We finally determined the proportionality relation between *p*_*att*_ and *Pe* from the slope of the curve, since *α*log(*x*) = log(*x*^*α*^).

